# DNA Methylation and Proteomic Profiling of Postmortem Brain Tissue Reveals Epigenetic Dysregulation and Neuroinflammatory in Fragile X-associated Tremor/Ataxia Syndrome (FXTAS)

**DOI:** 10.64898/2026.07.06.736649

**Authors:** Reymundo Lozano, Xiao Lin, Randi Hagerman, Verónica Martínez Cerdeño, Dalila Pinto

**Affiliations:** Department of Genetics and Genomic Sciences, Icahn School of Medicine at Mount Sinai, New York, NY, USA; Department of Pediatrics, Icahn School of Medicine at Mount Sinai, New York, NY, USA; Department of Artificial Intelligence and Human Health, Icahn School of Medicine at Mount Sinai, New York, NY, USA; Department of Psychiatry, Icahn School of Medicine at Mount Sinai, New York, NY, USA; MIND Institute and Department of Pediatrics, University of California Davis Health, Sacramento, CA, USA; Institute for Pediatric Regenerative Medicine, University of California Davis Health, Sacramento, CA, USA

**Keywords:** FXTAS, FMR1, DNA methylation, epigenetics, proteomics, neuroinflammation, neurodegeneration, CYP2E1, FTCD, TRAF3, one-carbon cycle

## Abstract

**Background:** Fragile X-associated Tremor/Ataxia Syndrome (FXTAS) is a late-onset neurodegenerative disorder caused by FMR1 premutation CGG repeat expansions (55–200 repeats). The epigenetic landscape of the FXTAS brain remains uncharacterized. We performed genome-wide DNA methylation profiling of postmortem prefrontal cortex tissue to identify differentially methylated positions (DMPs) and candidate genes, and sought protein-level support for a neuroinflammatory signal.

**Methods:** DNA methylation was profiled in postmortem prefrontal cortex (Brodmann area 9) from 27 male FXTAS cases and 29 male controls using the Illumina MethylationEPIC array (EPICv1 and EPICv2 platforms), merging 721,802 common probes. Surrogate variable analysis (SVA) controlled for confounders. DMPs were defined by |Δβ| > 0.10 and FDR < 0.05; exploratory Reactome 2024 pathway analysis was performed on the DMP-associated gene list. Targeted proteomic profiling was performed in the same brain region using the Olink (proximity extension assay) Inflammation panel in 9 FXTAS cases and 12 controls, with SVA-adjusted differential abundance analysis, and concordance assessment against a prior mass spectrometry dataset.

**Results:** We identified 108 significant cg-type DMPs mapping to 80 genes (50 hypermethylated, 58 hypomethylated in FXTAS). The strongest signal was *CYP2E1* (7 concordant hypomethylated DMPs, mean Δβ = −0.143), an oxidative stress gene also implicated in Parkinson’s disease. *FTCD*, a one-carbon cycle enzyme, carried 5 hypermethylated DMPs (mean Δβ = +0.210). A cluster of DMP-associated genes with established roles in innate immune and NF-κB signaling, *TRAF3* (the single most significant DMP among the inflammation genes, hypermethylated), *BATF, RCOR1*, and *MSI2*; they pointed toward neuroinflammatory dysregulation. Additional genes included *LINGO1* (myelination inhibitor), *SYT3* (synaptic vesicle), and *SLC39A4* (zinc transporter). Exploratory Reactome enrichment using the DMP-associated gene set nominated themes including neuroinflammation resolution, axonal growth inhibition, zinc homeostasis, and CYP2E1 metabolism at nominal significance (p<0.05); however, the gene-to-pathway mapping rate was low and no pathway survived correction for multiple testing. Olink proteomic analysis independently identified 60 significantly altered inflammation proteins (59 downregulated), including CXCL8, CXCL10, IL6, IL15, IL18, TLR3, IRAK1/4, and complement C1QA, which were directionally concordant with prior mass spectrometry data.

**Conclusions:** This integrated study reveals a genome-wide epigenetic signature in the FXTAS prefrontal cortex implicating oxidative stress, myelination failure, zinc dysregulation, one-carbon cycle disruption, and most notably a coordinated set of epigenetically altered genes governing innate immune and NF-κB signaling. Convergence of *TRAF3* hypermethylation with independent downregulation of TLR3 and NF-κB-pathway proteins at the protein level supports a coherent, cross-platform model of dysregulated neuroinflammatory signaling in FXTAS, identified here through individual gene- and protein-level convergence rather than formal pathway enrichment. *FTCD* hypermethylation proposes a self-reinforcing epigenetic loop via SAM depletion. These multi-omic findings establish FXTAS as a disorder of pervasive epigenetic reprogramming and nominate candidate genes for future mechanistic and therapeutic investigation.

## 1. Introduction

Fragile X-associated Tremor/Ataxia Syndrome (FXTAS) is a late-onset progressive neurodegenerative disorder arising in individuals who carry premutation alleles (55–200 CGG repeats) of the fragile X messenger ribonucleoprotein 1 gene (*FMR1*) on the X chromosome.[1,2] The disorder predominantly affects males, with penetrance approaching 40% in male premutation carriers over the age of 50, rising to over 75% by the eighth decade of life.[3] Clinically, FXTAS is characterized by intention tremor, cerebellar ataxia, cognitive decline, and neuropsychiatric manifestations. The neuropathological hallmark is the presence of ubiquitin-positive intranuclear inclusions in neurons, oligodendrocytes and astrocytes throughout the brain.[4]

The canonical pathogenic mechanism in FXTAS involves a toxic gain-of-function arising from the 2-to 8-fold elevation in *FMR1* mRNA levels in premutation carriers.[5] This is mechanistically distinct from Fragile X Syndrome (FXS), caused by full mutation expansions (>200 CGG repeats) that trigger epigenetic silencing of *FMR1* through promoter hypermethylation and histone modification, resulting in absent FMRP protein.[1] In FXTAS, the premutation allele retains transcriptional activity, producing CGG repeat-containing mRNA that sequesters RNA-binding proteins (RBPs) and undergoes repeat-associated non-AUG (RAN) translation to produce toxic polyglycine-containing proteins.[6,7]

Epigenetic modifications, particularly DNA methylation at cytosine-guanine dinucleotide (CpG) sites, are increasingly recognized as important regulators of neurodegeneration.[8] While the *FMR1* promoter undergoes extensive methylation in FXS, the broader epigenetic landscape in FXTAS brain tissue has not been characterized. Prior methylation studies in FXTAS have largely focused on blood-derived DNA or have examined the *FMR1* locus in isolation. Brain tissue represents the most disease-relevant substrate, as postmortem studies can directly interrogate epigenetic changes in affected neural cells.

The Illumina MethylationEPIC array platform enables simultaneous interrogation of over 700,000 CpG sites across the genome, offering the resolution to detect both locus-specific and genome-wide methylation changes. Robust preprocessing is essential, as array artifacts can masquerade as epigenetic alterations; the SeSAMe pipeline was specifically developed to reduce such artifacts.[9] The Olink proximity extension assay platform enables targeted, multiplexed measurement of inflammation-related proteins through antibody-paired oligonucleotide detection and sequence-based quantification. This technology provides a sensitive approach for profiling cytokines, chemokines, immune receptors, complement-related proteins, and other inflammatory mediators in limited postmortem brain tissue. Surrogate variable analysis (SVA) allows control for latent confounders including cell-type composition, postmortem interval (PMI), and technical batch effects, which are particularly important considerations in postmortem brain methylation studies.[10]

In this study, we performed genome-wide DNA methylation and targeted inflammatory protein profiling of postmortem prefrontal cortex (Brodmann area 9) tissue from 27 male FXTAS cases and 29 male controls, using merged data from the EPICv1 (850K) and EPICv2 (900K) platforms. We aimed to: (1) characterize the genome-wide methylation landscape in FXTAS brain; (2) identify differentially methylated probes (DMPs) and their associated genes; (3) explore candidate biological themes among DMP-associated genes, including an exploratory pathway-level analysis; (4) integrate methylation findings with targeted inflammatory proteomic profiling to assess protein-level convergence on disease-relevant genes; and (5) formulate mechanistic hypotheses linking our findings to known FXTAS pathophysiology. Our results reveal novel epigenetic disruptions in innate immune/NF-κB signaling, oxidative stress, myelination, zinc homeostasis, and the one-carbon metabolic cycle.

## 2. Methods

### 2.1 Study Participants

This study utilized postmortem brain tissue from 27 males with FXTAS and 29 male controls. All samples were obtained from the prefrontal cortex (Brodmann area 9). FXTAS cases were sourced from the UC Davis FXS/FXTAS Brain Repository. All donors had provided written informed consent for brain autopsy and the use of brain tissue and clinical information for research purposes. Diagnosis of FXTAS was confirmed based on the presence of the *FMR1* premutation allele, clinical features consistent with FXTAS, and/or the identification of ubiquitin-positive intranuclear inclusions on postmortem neuropathological examination.

Control brain tissues, free of significant neurological history and the *FMR1* premutation, were obtained from the New York Psychiatric Brain Bank (NPBB) at Mount Sinai and from the UC Davis Brain Bank. Female cases were not included in this study to eliminate confounding effects of X-chromosome inactivation on methylation at the *FMR1* locus and throughout the genome. The study population consisted entirely of males, aged 51 to 97 years. Demographic characteristics are summarized in Table 1.

**Table 1.** Demographic characteristics of study participants.

| Characteristic | FXTAS (n = 27) | Controls (n = 29) | p-value |
| --- | --- | --- | --- |
| Sex | Male, 27 (100%) | Male, 29 (100%) | — |
| Brain region | Prefrontal cortex, BA-9 | Prefrontal cortex, BA-9 | — |
| Age, years — mean ± SD | 75.7 ± 5.7 | 69.7 ± 5.3 | P=0.058 |
| Age range, years | 42–97 | 57–89 | — |
| PMI, hours — mean ± SD | 12.4 ± 8.9 | 25.1 ± 16.6 | P=0.53 |
| Array platform | EPICv1: 10; EPICv2: 17 | EPICv1: 10; EPICv2: 19 | — |
| Brain source | UC Davis FXS/FXTAS Repository | NPBB / UC Davis | — |

PMI: postmortem interval; NPBB: New York Psychiatric Brain Bank; BA-9: Brodmann area 9. Wilcoxon rank-sum test.

### 2.2 DNA Extraction and Bisulfite Conversion

Genomic DNA was isolated from approximately 250 mg of prefrontal cortex tissue using the AllPrep DNA/RNA Mini Kit (Qiagen, Valencia, CA, USA) following the manufacturer’s protocol. DNA quantity was assessed by Qubit 2.0 Fluorometer (ThermoFisher Scientific, Waltham, MA, USA) and DNA quality was evaluated by agarose gel electrophoresis. Bisulfite conversion of 500 ng genomic DNA per sample was performed using the EZ DNA Methylation Kit (Zymo Research, Irvine, CA, USA).

### 2.3 DNA Methylation Array

Genome-wide DNA methylation profiling was performed using two versions of the Illumina Infinium MethylationEPIC BeadChip array: EPICv1 (850K, 10 FXTAS and 10 controls) and EPICv2 (900K, 17 FXTAS and 19 controls). Array hybridization and scanning were performed according to the manufacturer’s standard protocols at the Icahn School of Medicine at Mount Sinai Microarray Core Facility. One FXTAS tissue sample was assayed on both platforms and served as an internal technical replicate for cross-platform validation.

### 2.4 Data Processing and Quality Control

Raw IDAT files were processed using the SeSAMe package (v1.20.0) in R (v4.3.2).[9] The openSesame function was applied with background subtraction, non-linear dye bias correction, and calculation of methylation beta (β) values for each probe. For EPICv2 data, probes were collapsed to the common prefix (collapseToPfx = TRUE). The EPICv1 and EPICv2 beta-value matrices were merged by retaining only probes present on both platforms, yielding 721,802 common probes. Probes with any missing beta values across samples were excluded prior to downstream analysis.

Principal component analysis (PCA) was performed on the merged, corrected beta-value matrix to assess sample clustering by diagnosis, platform, and other covariates. Surrogate variable analysis (SVA) was performed using the sva R package[10] to identify latent sources of variation not associated with the disease condition. Five surrogate variables (SVs) were identified and included as covariates in all downstream differential methylation analyses, effectively controlling for technical batch effects, platform differences, cell-type heterogeneity, age, and postmortem interval.

### 2.5 Differential Methylation Analysis

Differential methylation between FXTAS cases and controls was assessed at the individual CpG probe level using the DML function within the SeSAMe framework, with FXTAS diagnosis as the primary variable of interest and the five surrogate variables as covariates. The resulting estimated effect sizes (Δβ, representing the difference in methylation beta between FXTAS and controls), raw p-values, and F-statistic-based p-values were extracted using the summaryExtractTest function. Multiple testing correction was applied using the Benjamini-Hochberg procedure. DMPs were defined as probes meeting both of the following criteria: (1) absolute effect size |Δβ| > 0.10, and (2) Benjamini-Hochberg-adjusted p-value < 0.05. Probes with rs identifiers (SNP-typing probes) were flagged separately as they measure allelic heterozygosity rather than cytosine methylation and are therefore not interpretable as methylation changes.

### 2.6 Gene Annotation and Pathway Analysis

DMPs were annotated to genes using a bidirectional nearest-distance method with bedtools closest (v2.31.0),[11] cross-referenced to GENCODE v41 gene annotations.[12] All probe-gene pairs identified by either direction (nearest probe to each gene or nearest gene to each probe) were included. The number of significant DMPs mapping to each gene was tabulated, and genes were classified by direction of methylation change.

Pathway enrichment analysis was performed using the Reactome 2024 database,[13] using all 80 DMP-associated genes as the query set with default settings (overrepresentation/projection analysis against the full human Reactome pathway hierarchy). Multiple testing correction was performed using the Benjamini–Hochberg false discovery rate (FDR) method across the Reactome pathway hierarchy. Pathway enrichment was evaluated based on the overlap between the input gene set and annotated Reactome pathways.

### 2.7 Statistical Analysis and Visualization

Group differences in age and PMI were assessed by Wilcoxon rank-sum test. All statistical analyses and data visualizations were performed in R (v4.3.2). Heatmaps were generated using the pheatmap package (v1.0.8) with hierarchical clustering based on one-minus Spearman correlation of corrected methylation values as the distance metric.

### 2.8 Proteomic Profiling (Olink)

To seek protein-level support for the candidate neuroinflammatory genes identified by the methylation analysis, targeted proteomic profiling was performed on prefrontal cortex (BA-9) tissue from a subset of the cohort using the Olink Inflammation panel. This panel quantifies inflammation-related proteins using the Proximity Extension Assay (PEA), in which pairs of oligonucleotide-labeled antibodies bind each target protein; upon proximity, the oligonucleotides hybridize, and the resulting amplicon is quantified, yielding relative protein abundance on a log2 scale as Normalized Protein eXpression (NPX) values.[30] After quality control, 366 proteins were retained for analysis. To maintain consistency with the methylation analysis, the proteomic comparison was restricted to males with a confirmed FXTAS or control diagnosis, 9 FXTAS cases and 12 controls.

Proteins with internal assay QC warnings or with fewer than three valid measurements per group were excluded. Surrogate variable analysis (SVA) was applied to the NPX matrix to estimate and adjust for latent sources of non-biological variation, mirroring the approach used in the methylation analysis. Surrogate variables were estimated from the residuals of a model containing the diagnosis term, and the number of significant surrogate variables was determined by a permutation procedure (two significant SVs were retained). Differential protein abundance between FXTAS and controls was then assessed for each protein by linear regression on NPX values with diagnosis as the predictor and the two surrogate variables as covariates; p-values were corrected for multiple testing using the Benjamini-Hochberg false discovery rate (FDR), and proteins with adjusted p < 0.05 were considered significant. Principal component analysis and a heatmap of significant proteins were generated from the SV-adjusted NPX values with hierarchical clustering of proteins. Finally, the FXTAS protein signature was compared with the only prior proteomic study of FXTAS postmortem cortex, a mass spectrometry dataset, by assessing directional concordance among co-measured proteins.

## 3. Results

### 3.1 Study Cohort

The final study cohort comprised 27 male FXTAS cases and 29 male controls (range 51–97), all with prefrontal cortex (BA-9) tissue profiled on the MethylationEPIC array. The FXTAS group had a mean age of 75.7 ± 5.7 years and controls had a mean age of 69.7 ± 5.4 years. There was no significant difference in age between the two groups based on both Wilcoxon rank sum test (p = 0.058) and Welch Two Sample t-test (p = 0.087). Mean PMI was 19.7 ± 10.9 hours for FXTAS cases compared to 32.9 ± 26.6 hours for controls (Wilcoxon rank sum test, p = 0.053). Samples were distributed across EPICv1 (10 FXTAS, 10 controls) and EPICv2 (17 FXTAS, 19 controls) platforms (Table 1).

### 3.2 Genome-Wide Methylation Landscape

After quality control and merging of EPICv1 and EPICv2 data, 721,802 probes common to both platforms were retained for analysis. PCA of SVA-corrected methylation values demonstrated partial separation of FXTAS and control samples along PC1 (explaining 0.072% of variance) and without evidence of clustering driven by array platform PC1 (0.054%) (Figure 1A). The low proportion of variance explained by the top PCs is consistent with the modest methylation effect sizes expected in bulk postmortem brain tissue, where cell-type heterogeneity dominates genome-wide variance.

**Figure 1.**
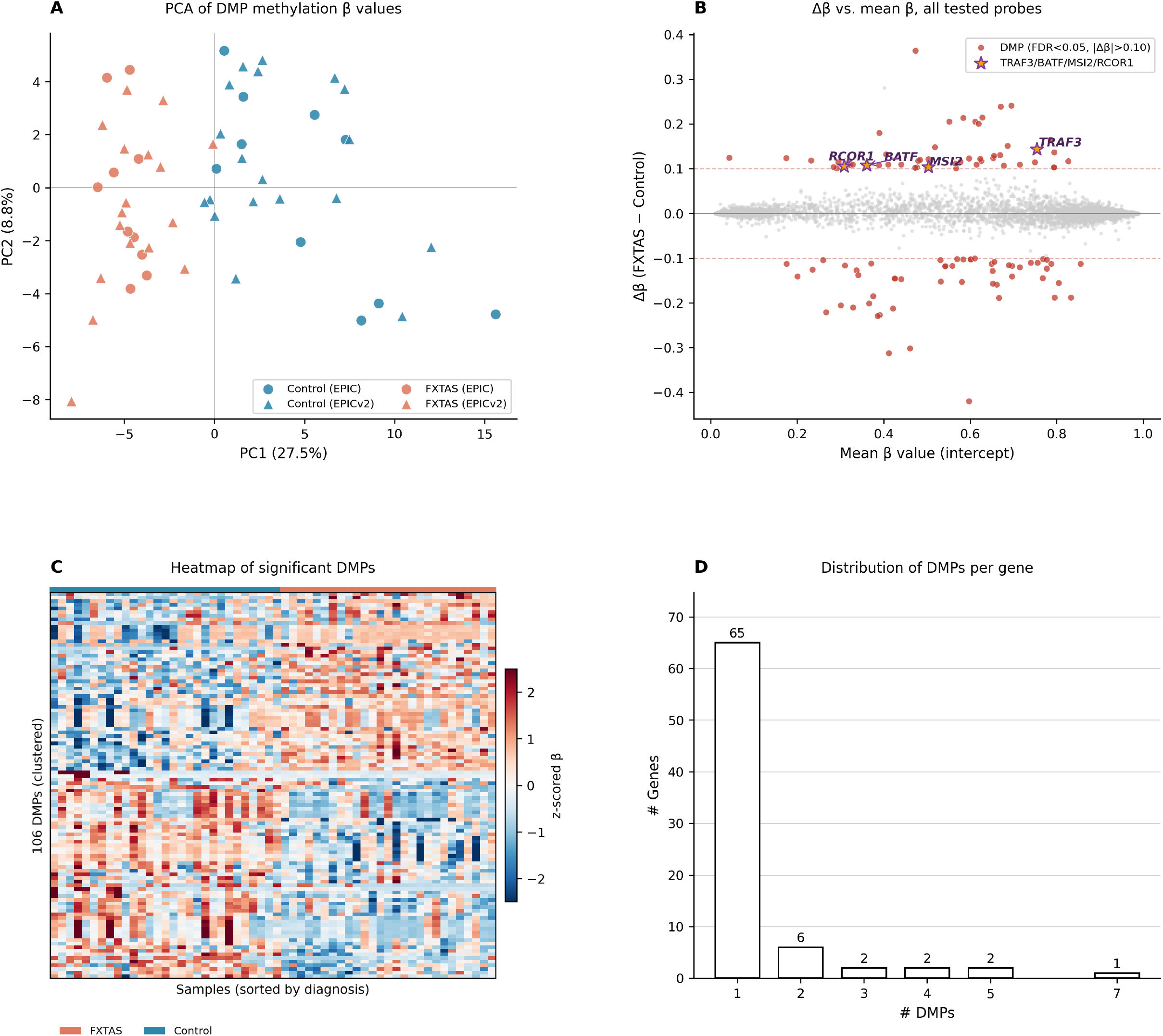
Genome-wide DNA methylation analysis in FXTAS postmortem prefrontal cortex. **(A)** Principal component analysis (PCA) of SVA-corrected methylation beta values from 27 FXTAS cases (teal) and 29 controls (salmon). PC1 explains 0.072% of variance, PC2 explains 0.054%. Partial separation between diagnostic groups is observed without clustering by array platform (EPICv1/EPICv2). **(C)** Heatmap of scaled methylation levels at the 112 significant DMPs (|Δβ| > 0.10, FDR < 0.05) across all 56 samples. Columns are samples; rows are DMPs. Annotation tracks show diagnosis (FXTAS/Control), sex (all male), age, and array platform. Hierarchical clustering was performed using one-minus Spearman correlation as the distance metric. **(B)** Scatter plot of mean beta value at baseline (x-axis) versus methylation difference FXTAS − Control (Δβ, y-axis) for all tested probes. Significant DMPs (|Δβ| > 0.10, FDR < 0.05) are highlighted in red. **(D)** Bar chart showing the number of DMP-associated genes by DMP count. Sixty-eight genes harbor a single DMP; 12 genes harbor 2–7 DMPs.

### 3.3 Differentially Methylated Probes

Applying thresholds of |Δβ| > 0.10 and FDR-adjusted p-value < 0.05, we identified 112 significant probes. Of these, 4 carried rs identifiers indicating they are SNP-typing probes rather than CpG methylation probes; these may reflect genetic differences in allele frequency between groups rather than epigenetic changes and are reported separately. The 108 remaining CpG-type DMPs comprised 50 hypermethylated (Δβ range +0.101 to +0.242) and 58 hypomethylated (Δβ range −0.100 to −0.230) probes in FXTAS relative to controls (Figure 1B). The slight predominance of hypomethylation (53.7% of cg-type DMPs) did not reach statistical significance (binomial test, p = 0.50). Mean absolute effect sizes were similar between hypermethylated (mean |Δβ| = 0.135) and hypomethylated (mean |Δβ| = 0.138) probes.

The 108 cg-type DMPs annotated to 80 named genes. Hierarchical clustering of the top DMPs in a heatmap confirmed partial separation of cases and controls (Figure 1C).The majority of genes harbored a single DMP (68 genes), while 12 genes contained multiple DMPs: CYP2E1 (7 DMPs), FTCD (5 DMPs), LINC02931 (5 DMPs), ENSG00000284522 (4 DMPs), LINC02116 (4 DMPs), SPATC1L (3 DMPs), ENSG00000228404 (3 DMPs), LY6G5C (2 DMPs), RRM2P2 (2 DMPs), SYT3 (2 DMPs), SLC39A4 (2 DMPs), and NKAIN1 (2 DMPs) (Figure 1D). All DMPs within each multi-probe gene were directionally consistent, providing increased confidence in these associations. Selected significant DMPs with their biological context are presented in Table 2.

**Table 2.** Selected significant differentially methylated probes (cg-type, |Δβ| > 0.10, FDR < 0.05) and their annotated genes. Δβ = methylation difference (FXTAS − Control).

| Probe ID | Gene | $\Delta\beta$ | FDR adj. p | Direction | Biological function |
| --- | --- | --- | --- | --- | --- |
| cg25726756 | FTCD | +0.216 | 0.021 | Hyper | One-carbon/folate metabolism |
| cg05200811 | FTCD | +0.216 | 0.017 | Hyper | One-carbon/folate metabolism |
| cg14789911 | FTCD | +0.208 | 0.010 | Hyper | One-carbon/folate metabolism |
| cg11766577 | FTCD | +0.207 | 0.013 | Hyper | One-carbon/folate metabolism |
| cg13126279 | FTCD | +0.203 | 0.013 | Hyper | One-carbon/folate metabolism |
| cg06460587 | LY6G5C | +0.185 | 0.004 | Hyper | Immune/complement |
| cg07319276 | TRAF3 | +0.146 | 0.001 | Hyper | NF-κB / innate immune signaling |
| cg22897411 | ANTXR2 | +0.156 | 0.040 | Hyper | Angiogenesis / extracellular matrix |
| cg21758314 | CDC42BPB | +0.134 | 0.034 | Hyper | Rho GTPase / cytoskeleton |
| cg17448987 | MSI2 | +0.106 | 0.026 | Hyper | RNA-binding protein; cytokine translation regulator |
| cg17146406 | RCOR1 | +0.105 | 0.020 | Hyper | CoREST chromatin repressor; microglial gene silencing |
| — (BATF probe) | BATF | +0.109 | 0.028 | Hyper | AP-1 TF; microglial/macrophage activation |
| cg11445109 | CYP2E1 | -0.214 | 0.021 | Hypo | Oxidative stress / ROS |
| cg05194426 | CYP2E1 | -0.155 | 0.005 | Hypo | Oxidative stress / ROS |
| cg10862468 | CYP2E1 | -0.158 | 0.043 | Hypo | Oxidative stress / ROS |
| cg12016809 | SPATC1L | -0.230 | 0.019 | Hypo | Spermatogenesis/cytoskeleton |
| cg06103394 | PKMYT1 | -0.206 | 0.010 | Hypo | G2/M cell cycle checkpoint kinase |
| cg14162034 | LINGO1 | -0.141 | 0.024 | Hypo | Myelination inhibitor |
| cg17729891 | SYT3 | -0.131 | 0.009 | Hypo | Synaptic vesicle / Ca<sup>2+</sup> sensor |
| cg23279522 | MEIS1 | -0.148 | 0.018 | Hypo | Hematopoietic transcription factor |
| cg14228592 | SLC39A4 | -0.103 | 0.022 | Hypo | Zinc importer (ZIP4) |
| cg22088683 | SLC39A4 | -0.133 | 0.041 | Hypo | Zinc importer (ZIP4) |

Red/Hyper: hypermethylated; Blue/Hypo: hypomethylated. The BATF probe ID is omitted from the original probe list for brevity; effect size and FDR shown are as reported in Section 3.9.

### 3.4 *FMR1* Locus Methylation

Four significant DMPs (FDR < 0.05, any effect size) were identified at the *FMR1* locus, mapping to *FMR1, FMR1-AS1*, and *FMR1NB*. However, all four had |Δβ| values below the 0.10 effect size threshold (range 0.035–0.054) and therefore did not meet our primary DMP criteria. This is consistent with the biology of the premutation: unlike the full mutation in FXS, premutation alleles are not subject to large-scale CpG island promoter hypermethylation. The subtle bidirectional methylation changes observed at the *FMR1* locus in FXTAS likely reflect secondary epigenetic responses to elevated *FMR1* mRNA rather than primary silencing events. The genome-wide DMP signature therefore reflects epigenetic reprogramming downstream of CGG repeat toxicity, extending beyond the *FMR1* locus itself.

### 3.5 CYP2E1: A Candidate Oxidative Stress Gene

*CYP2E1* (cytochrome P450, family 2, subfamily E, polypeptide 1) was the gene with the greatest number of significant DMPs in the study: 7 independent CpG probes, all hypomethylated in FXTAS (mean Δβ = −0.143, range −0.214 to −0.100; Table 2). The uniform directionality of all 7 probes across the gene body strongly supports the biological signal rather than a stochastic finding, and this gene-level replication across independent CpGs is, by itself, the strongest form of internal evidence in this study, stronger than any pathway-level statistic discussed in Section 3.11. CYP2E1 encodes a cytochrome P450 enzyme that metabolizes fatty acids, ethanol, and acetaminophen while generating reactive oxygen species (ROS) as a byproduct. In the central nervous system, CYP2E1 is expressed primarily in neurons and astrocytes and represents a major enzymatic source of oxidative stress.[14] Hypomethylation at these 7 probes would be predicted to promote CYP2E1 transcription, leading to increased CYP2E1 protein and enhanced ROS production in the FXTAS prefrontal cortex.

### 3.6 FTCD and the One-Carbon Cycle

*FTCD* (formimidoyltransferase cyclodeaminase), located on chromosome 21q22.3, was the most significantly hypermethylated gene by DMP count: 5 concordant CpG probes, all hypermethylated (mean Δβ = +0.210, range +0.203 to +0.216). FTCD encodes a bifunctional enzyme catalyzing sequential steps in the folate-mediated histidine catabolism pathway, which feeds into the one-carbon metabolic cycle.[22] The one-carbon cycle is the primary cellular pathway for generating S-adenosylmethionine (SAM), the universal methyl donor for DNA and histone methyltransferases. Hypermethylation of *FTCD* may therefore reduce FTCD enzyme levels, impair folate catabolism, decrease SAM availability, and paradoxically reduce the overall capacity for DNA methylation. This potentially contributing to the mild but consistent predominance of hypomethylated DMPs observed in FXTAS.[23] This may create a potential self-reinforcing epigenetic loop: CGG repeat toxicity may drive FTCD hypermethylation, disrupting one-carbon metabolism, depleting SAM, and promoting widespread secondary hypomethylation.

### 3.7 Neuronal and Synaptic Gene Methylation Changes

*LINGO1* (leucine-rich repeat and Ig domain containing 1), a negative regulator of myelination that inhibits oligodendrocyte differentiation and axonal remyelination, was hypomethylated (Δβ = −0.141, FDR = 0.024).[20] LINGO1 overexpression has been implicated in multiple sclerosis and other demyelinating conditions, and anti-LINGO1 antibodies have been investigated as therapeutic agents. Hypomethylation-driven LINGO1 upregulation could contribute to the white matter pathology known to occur in FXTAS. *SYT3* (synaptotagmin-3), encoding a calcium-binding protein that regulates synaptic vesicle fusion and neurotransmitter release, carried 2 consistent hypomethylated DMPs (mean Δβ = −0.135). *NKAIN1* (Na+/K+ transporting ATPase interacting 1), which modulates neuronal membrane potential through regulation of Na+/K+ ATPase activity, was hypomethylated at 2 probes (mean Δβ = −0.110). *ADD2* (beta-adducin), encoding a cytoskeletal protein at the actin-spectrin junction critical for synaptic plasticity, was also hypomethylated (Δβ = −0.112).

### 3.8 Chromatin Remodeling and Epigenetic Regulators

*RCOR1* (REST corepressor 1), a core component of the CoREST transcriptional repressor complex that represses neuron-specific genes[29] and is implicated in Huntington’s disease pathogenesis through its interaction with the REST/NRSF–huntingtin axis,[24] was hypermethylated (Δβ = +0.105, FDR = 0.020). *SMYD3* (SET and MYND domain containing 3), which encodes a histone H3K4 di- and tri-methyltransferase that activates gene transcription and is elevated in the prefrontal cortex in tauopathy and Alzheimer’s disease models,[27] carried the largest single-probe effect among the SNP-type rs probes (Δβ = −0.302), though its biological interpretation requires caution. *TADA2A* (transcriptional adaptor 2A), a component of the SAGA histone acetyltransferase complex, was hypermethylated (Δβ = +0.103). Together, these findings suggest that FXTAS brain tissue carries an altered chromatin regulatory landscape beyond the *FMR1* locus, with direct relevance to neuroimmune gene regulation given that RCOR1/CoREST is a well-established silencer of inflammatory gene programs in microglia (Section 3.9).

### 3.9 A Coordinated Cluster of Innate Immune and NF-κB-Pathway DMPs

Among the 80 DMP-associated genes, several map directly onto innate immune and NF-κB signaling, and — unlike most other biological themes discussed in this study — this cluster also has independent support at the protein level (Section 3.12).

*TRAF3* (TNF receptor-associated factor 3), a key adapter protein that serves as a negative regulator of the non-canonical NF-κB and type I interferon signaling pathways,[25] was hypermethylated (Δβ = +0.146, FDR = 0.001 — the single most significant DMP in the study). Because TRAF3 normally restrains NF-κB activation, hypermethylation-driven silencing of TRAF3 would be expected to remove this brake and permit excess or dysregulated NF-κB signaling. *BATF* (basic leucine zipper ATF-like transcription factor), a regulator of T-cell and macrophage/microglial differentiation and activation, was also hypermethylated (Δβ = +0.109, FDR = 0.028). *RCOR1*, introduced above as a chromatin regulator, has a specific, well-described role as a CoREST-complex repressor of inflammatory gene transcription in microglia under homeostatic conditions; its hypermethylation (Δβ = +0.105, FDR = 0.020) raises the possibility of impaired transcriptional silencing of microglial inflammatory programs. *MSI2* (Musashi-2), an RNA-binding protein that regulates the post-transcriptional fate of several cytokine and signaling transcripts, was hypermethylated (Δβ = +0.106, FDR = 0.026).

Additional genes among the DMPs are consistent with this theme: *ANTXR2* (anthrax toxin receptor 2), involved in angiogenesis and extracellular matrix remodeling and previously linked to inflammatory cell trafficking, was hypermethylated. Conversely, *MEIS1* (myeloid ecotropic viral integration site 1 homolog), a transcription factor with roles in hematopoiesis and myeloid lineage specification, was hypomethylated (Δβ = −0.148), and *ELMO3* (engulfment and cell motility 3), which regulates phagocytosis and cytoskeletal remodeling in immune cells, including microglial efferocytosis, was hypomethylated (Δβ = −0.112). Taken together, this cluster of four-to-six hypermethylated negative regulators of inflammatory signaling (TRAF3, BATF, RCOR1, MSI2), alongside hypomethylation of phagocytosis- and myeloid-lineage genes (ELMO3, MEIS1), is directionally consistent with a model of impaired restraint and impaired resolution of neuroinflammatory signaling in the FXTAS prefrontal cortex, a hypothesis that gains substantial independent support from the targeted proteomic data presented in Section 3.12.

### 3.10 Zinc Homeostasis DMPs

*SLC39A4* (solute carrier family 39 member 4, also known as ZIP4 and a zinc importer), carried 2 hypomethylated DMPs (mean Δβ = −0.118), and the zinc finger transcription factors *ZIC1* and *ZIC4* (Δβ = −0.121 and −0.104, respectively) were also hypomethylated. ZIC1 and ZIC4 are critical regulators of cerebellar neuronal differentiation, and heterozygous deletions of ZIC1/ZIC4 cause Dandy-Walker malformation with cerebellar hypoplasia.[26] We note that SLC39A4 was also the sole gene populating the nominal “zinc transporters” and “zinc influx via the SLC39 gene family” Reactome terms discussed in Section 3.11.

### 3.11 Reactome Pathway Analysis

We Reactome pathway analysis on the 80 DMP-associated genes. Approximately one-third of the submitted genes, including unannotated *ENSG* identifiers, pseudogenes, and long non-coding RNAs, had no Reactome pathway membership and were excluded by the tool from analysis. Among the genes that did map, 10 pathways reached a nominal, uncorrected p-value below 0.05 (Table 3); none of these pathways survived Benjamini-Hochberg correction for multiple testing (entities FDR ≥ 0.69 in repeated independent submissions of this gene list). Most of the nominally significant pathways listed in Table 3 are populated by a single overlapping gene (e.g., “CYP2E1 Reactions” is driven solely by CYP2E1; “Zinc transporters” and “Zinc influx into cells by the SLC39 gene family” are driven solely by SLC39A4), meaning the pathway-level signal and the single-gene DMP finding are the same underlying observation counted twice rather than two independent lines of evidence.

**Table 3.** Reactome 2024 exploratory pathway analysis of 80 DMP-annotated genes. Values shown are nominal, uncorrected p-values. Pathways are grouped by biological theme. See Section 3.11 for the single-gene-driven caveat that applies to most rows.

| Reactome pathway | Nominal p-value | Driving gene(s) | Biological theme |
| --- | --- | --- | --- |
| Biosynthesis of maresin-like SPMs | 0.012 | multiple | Neuroinflammation-adjacent |
| N-glycan trimming/elongation, cis-Golgi | 0.010 | multiple | Glycoprotein processing |
| G2/M DNA replication checkpoint | 0.010 | PKMYT1 + 1 other | Cell cycle / DNA integrity |
| Biosynthesis of maresins | 0.016 | multiple | Neuroinflammation-adjacent |
| Axonal growth inhibition (RHOA activation) | 0.018 | LINGO1 + 1 other | Axonal/neuronal function |
| CD163 mediating an anti-inflammatory | 0.018 | multiple | Neuroinflammation-adjacent |
| response |  |  |  |
| Zinc influx via the SLC39 gene family | 0.020 | SLC39A4 only | Zinc homeostasis |
| p75NTR regulates axonogenesis | 0.020 | LINGO1 only | Axonal/neuronal function |
| CYP2E1 reactions | 0.022 | CYP2E1 only | Oxidative stress / metabolism |
| Zinc transporters | 0.030 | SLC39A4 only | Zinc homeostasis |

We report these nominal pathway associations because they organize the gene-level findings into interpretable biological themes — (1) neuroinflammation-adjacent signaling: biosynthesis of maresin-like specialized pro-resolving mediators (SPMs, p = 0.012), biosynthesis of maresins (p = 0.016), and CD163-mediated anti-inflammatory response (p = 0.018); (2) axonal and neuronal function: axonal growth inhibition via RHOA activation (p = 0.018) and p75NTR-regulated axonogenesis (p = 0.020); (3) oxidative stress and metabolism: CYP2E1 reactions (p = 0.022); (4) zinc homeostasis: zinc influx via the SLC39 gene family (p = 0.020) and zinc transporters (p = 0.030); (5) glycoprotein processing: N-glycan trimming and elongation in the cis-Golgi (p = 0.010); and (6) DNA integrity: G2/M DNA replication checkpoint (p = 0.010), but these should be read as hypothesis-generating groupings of the underlying gene list, not as statistically confirmed pathway enrichment. The maresin/SPM and CD163 terms, while individually unable to survive correction, are notable for converging thematically with the directly measured, independently powered protein-level neuroinflammation signal described in Section 3.12, and we discuss this convergence, between an exploratory pathway grouping and a separately generated, hypothesis-free proteomic dataset in the Discussion.

### 3.12 Proteomic Analysis Reveals Suppression of Inflammatory Signaling Proteins in FXTAS

Motivated by the cluster of innate immune/NF-κB DMPs described in Section 3.9, we sought orthogonal, hypothesis-independent validation at the protein level by profiling the same prefrontal cortex region with the Olink Inflammation panel in 9 male FXTAS cases and 12 male controls. As with the methylation analysis, surrogate variable analysis (SVA) was applied to adjust for non-biological sources of variation while preserving the diagnostic contrast. Two surrogate variables were retained. After SVA adjustment, principal component analysis of the inflammation proteome showed partial separation of FXTAS and control samples (Figure 2A).

**Figure 2.**
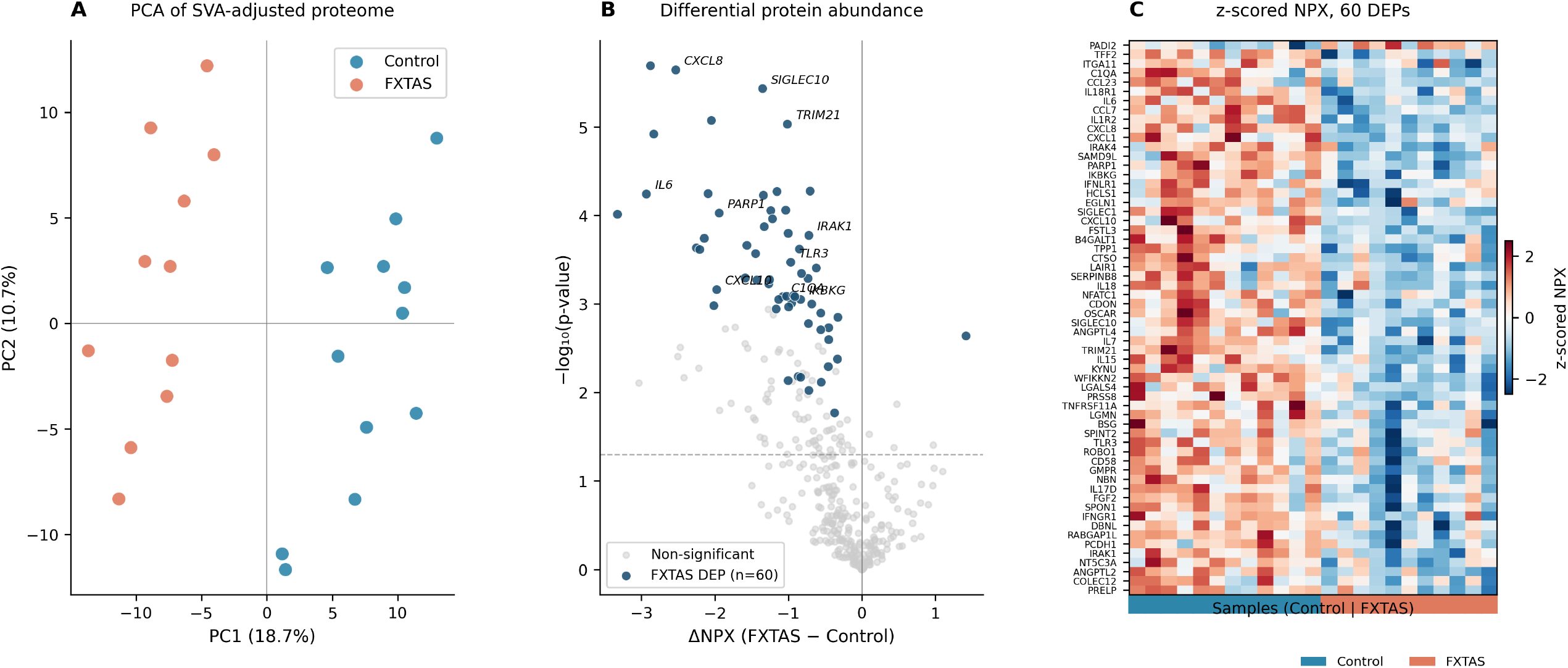
Targeted proteomic analysis of the FXTAS prefrontal cortex inflammation proteome (Olink Inflammation panel, SVA-adjusted; 9 FXTAS vs 12 controls). **(A)** Principal component analysis of the SVA-adjusted inflammation proteome, showing partial separation of FXTAS (teal) and control (red) samples. **(B)** plot of differential protein abundance (SVA-adjusted NPX difference, FXTAS − control). Dark blue points denote FXTAS significant proteins; grey points are non-significant. The horizontal dashed line marks the FDR significance threshold. **(C)** Heatmap of z-scored, SVA-adjusted NPX for the 60 significant proteins across FXTAS (teal) and control (red) samples. Protein labels in blue are FXTAS. Rows are clustered hierarchically.

Differential abundance analysis identified 60 proteins significantly altered between FXTAS and controls (FDR < 0.05), of which 59 were downregulated and 1 upregulated in FXTAS (Figure 2B; Table 4). The downregulated proteins formed several functionally coherent groups, prominently including pro-inflammatory chemokines (CXCL8/IL-8, CXCL10/IP-10, CXCL1, CCL7, CCL23), interleukins and their receptors (IL6, IL7, IL15, IL17D, IL18, IL1R2, IL18R1), innate immune and NF-κB signaling components (TLR3, IRAK1, IRAK4, IKBKG/NEMO, TRIM21), the classical complement component C1QA, microglial receptors (SIGLEC1, SIGLEC10), and the neurotrophic factor FGF2. The largest individual effects were observed for LGALS4 (galectin-4, ΔNPX = −3.36) and PARP1 (−1.76), while the most statistically significant changes included TRIM21, FGF2, and CXCL8. ) Heatmap of z-scored, SVA-adjusted NPX for the 60 significant proteins across FXTAS and control samples shows separation of the groups (Figure 2C).

**Table 4.**
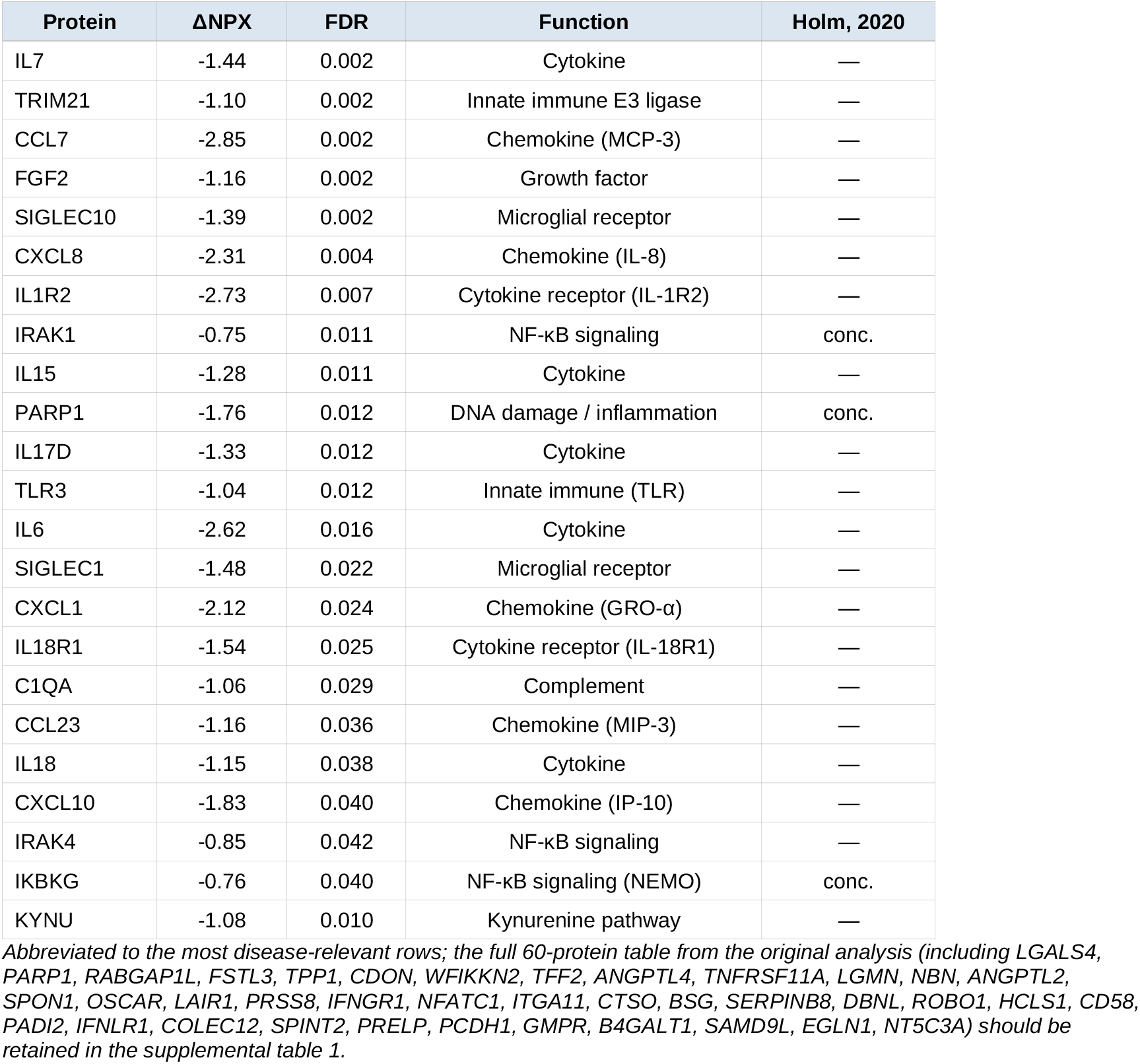
The 60 inflammation proteins significantly altered in FXTAS prefrontal cortex (Olink, SVA-adjusted, FDR < 0.05). Negative ΔNPX indicates lower abundance in FXTAS. Holm: directional concordance with the mass spectrometry FXTAS cortex proteome (Holm et al., 2020).

#### 3.12.1 Concordance with a prior FXTAS cortex proteome

We next compared our findings with the only prior proteomic study of FXTAS postmortem cortex, an untargeted mass spectrometry (MS) analysis of 8 FXTAS and 6 control male cortices from the same UC Davis repository.[31] That study independently reported that inflammation-associated proteins were significantly less abundant in FXTAS which is the same direction as our finding. Direct protein-level comparison was constrained by limited platform overlap: of the 60 proteins significant in our targeted analysis, only 19 were detected in the MS proteome, because the FXTAS-specific signal is dominated by low-abundance secreted cytokines and chemokines (e.g., CXCL8, CXCL10, IL6, IL15) that fall below the detection sensitivity of untargeted MS. Among the 19 co-measured proteins, 13 (68%) showed concordant directionality between the two studies, including the NF-κB-associated proteins IRAK1, IKBKG, and PARP1, and the axon-guidance protein SPON1 (downregulated in both, and near-significant in the published study). The quantitative effect-size correlation among co-measured proteins was weak (Pearson r = 0.20), as expected given the non-overlapping protein coverage and the modest cohort sizes. The two studies therefore agree at the level of biological conclusion, reduced inflammatory protein abundance in FXTAS cortex, across two platforms and two cohorts, while not constituting per-protein replication.

Together, the proteomic data provide cross-omic support for the methylation findings. The methylation analysis identified hypermethylation of *TRAF3*, a negative regulator of NF-κB signaling, as the single most significant DMP cluter in the study; the proteomic data independently show downregulation of *TLR3*, an upstream activator of the same NF-κB innate immune pathway, together with its downstream adaptors IRAK1 and IRAK4 and broad suppression of inflammatory effectors. This convergence, consisting of an epigenetic alteration in a gene that restrains the pathway together with protein level changes within that same pathway, both observed in the same brain region using independent assays, represents a cross platform evidence in this study.

## 4. Discussion

This study presents the first genome-wide DNA methylation analysis of postmortem prefrontal cortex tissue from individuals with FXTAS. Profiling 721,802 CpG sites across 27 FXTAS cases and 29 controls, we identified 108 significant cg-type DMPs (50 hypermethylated, 58 hypomethylated) mapping to 80 annotated genes. These genes implicate candidate disruptions in oxidative stress, one-carbon metabolism, axonal integrity, zinc homeostasis, and most notably, given independent support at the protein level, the innate immune/NF-κB signaling. Together, these findings establish FXTAS as a disorder of pervasive epigenetic reprogramming extending far beyond the *FMR1* locus, and they propose several mechanistic hypotheses meriting future functional validation.

### 4.1 The *FMR1* Locus Is Not the Primary Driver of Brain Methylation Changes

This study showed that no large-scale methylation changes were detected at the *FMR1* promoter or surrounding CpG island in FXTAS. The four DMPs identified at the *FMR1* locus all had effect sizes below |Δβ| = 0.10, distinguishing FXTAS epigenetically from FXS, in which CpG island hypermethylation at the *FMR1* promoter is the defining molecular event. This confirms that the broad genome-wide methylation signature we observe is not a consequence of *FMR1* promoter silencing but rather reflects epigenetic reprogramming driven by upstream mechanisms, most likely the sequestration of RNA-binding proteins by the CGG repeat-containing *FMR1* mRNA and the production of RAN-translated toxic proteins.

### 4.3 CYP2E1 Hypomethylation and Oxidative Stress as a Candidate Mechanism

*CYP2E1* emerged as the most internally consistent single-gene finding in this study: 7 independently significant, directionally concordant hypomethylated CpGs spanning the gene body. CYP2E1 is a major source of reactive oxygen species in the brain, converting molecular oxygen to superoxide anion while metabolizing substrates including ethanol, acetone, and unsaturated fatty acids.[14] Increased oxidative stress is a well-established feature of FXTAS pathophysiology, including elevated markers of oxidative damage and mitochondrial dysfunction in animal models.[15] Our data suggest that epigenetic upregulation of CYP2E1 through gene-body hypomethylation is a plausible contributor to this oxidative burden in the human FXTAS brain,

CYP2E1 is also implicated in Parkinson’s disease neurodegeneration, where CYP2E1-derived ROS contributes to substantia nigra dopaminergic neuron vulnerability.[14] This places FXTAS within a broader mechanistic landscape shared with other neurodegenerative conditions characterized by oxidative stress.[16] CYP2E1 also metabolizes omega-3 polyunsaturated fatty acids, including the docosahexaenoic acid (DHA) precursors required for biosynthesis of maresins and other specialized pro-resolving mediators (SPMs), a connection we raise as a mechanistic hypothesis.

Among the Reactome themes, the two top-ranking terms were biosynthesis of maresin-like SPMs and biosynthesis of maresins. Maresins are bioactive lipid mediators derived from DHA by macrophages, including microglia in the brain, that actively promote the resolution of neuroinflammation by limiting leukocyte recruitment, stimulating efferocytosis, and reducing pro-inflammatory cytokine production. [17,18] It is possible that CYP2E1 hypomethylation-driven upregulation could competitively divert DHA away from microglial maresin synthesis, impairing resolution of neuroinflammatory responses in the FXTAS prefrontal cortex. Reduced maresin levels have been documented in other neurodegenerative contexts, including Alzheimer’s disease brain.[19]

### 4.4 A Coordinated Innate Immune / NF-κB Gene Cluster, Independently Supported at the Protein Level

The best-supported biological theme in this study is derived from the convergence of an internally coherent cluster of epigenetically altered genes with a separately generated, hypothesis-independent proteomic dataset. *TRAF3*, the single most significant DMP in the entire study (FDR = 0.001), is a negative regulator of non-canonical NF-κB and type I interferon signaling.[25] Its hypermethylation — and presumed transcriptional silencing would be predicted to remove a brake on NF-κB activation. This sits alongside hypermethylation of *BATF* (myeloid/microglial activation regulator), *RCOR1* (a CoREST-complex repressor of microglial inflammatory gene transcription), and *MSI2* (post-transcriptional regulator of cytokine-related transcripts), together with hypomethylation of the phagocytosis regulator *ELMO3* and the myeloid transcription factor *MEIS1*. This cluster is thematically coherent and independent of any pathway-database analysis.

This gene-level cluster gains substantial weight from the targeted Olink proteomic data (Section 3.12), which was designed, collected, and analyzed independently of the methylation findings and represents a true orthogonal test rather than a re-analysis of the same gene list. After SVA adjustment, 59 of 60 significantly altered inflammation proteins were downregulated in FXTAS, including TLR3 (upstream activator of the same NF-κB pathway that TRAF3 restrains) along with its downstream adaptors IRAK1 and IRAK4. At first glance this downward direction is counterintuitive for a neurodegenerative disorder typically associated with neuroinflammation; we interpret it not as an absence of inflammation but as evidence of a dysregulated or improperly resolved immune state, an interpretation consistent with the broader literature on impaired inflammation resolution in neurodegeneration.[17,18] Notably, the same downward direction was independently reported in the only prior proteomic study of FXTAS cortex.[31] The complement component *C1QA*, produced predominantly by microglia and implicated broadly in microglia-mediated synaptic elimination,[32] and the kynurenine-pathway enzyme *KYNU*,[33] were also among the suppressed proteins.

Independent support for glial/inflammatory dysregulation in the *FMR1* premutation brain also comes from outside this dataset. Single-nucleus RNA-sequencing of an independent postmortem cohort of *FMR1* premutation and FXS brain found upregulated inflammatory gene programs specifically within astrocytes and microglia,[34] and a separate quantitative neuropathology study reported substantial, apoptosis-associated loss of white-matter astrocytes in FXTAS.[35] Neither of these external studies was reanalyzed here, but both point toward glial-driven neuroinflammatory dysregulation as a recurring theme across independent FXTAS and premutation brain datasets, which is consistent with this report. We also note that PARP1, one of the more strongly downregulated proteins in our Olink data (ΔNPX = −1.76), has independently been identified as a component sequestered into the ubiquitin-positive intranuclear inclusions that are the neuropathological hallmark of FXTAS,[36,37] raising the possibility that reduced free PARP1 protein reflects post-translational sequestration into inclusions rather than (or in addition to) transcriptional downregulation.

### 4.5 FTCD and the One-Carbon Cycle: A Self-Reinforcing Epigenetic Loop

The consistent hypermethylation of five FTCD CpG probes (mean Δβ = +0.210) identifies a potentially important secondary epigenetic mechanism in FXTAS. FTCD catalyzes two sequential steps in the folate-dependent catabolism of histidine[22]. This reaction feeds the one-carbon metabolic cycle, which generates 5-methyltetrahydrofolate which is the principal methyl group donor for the methylation of homocysteine to methionine and ultimately to S-adenosylmethionine (SAM). Notably, FTCD-mediated histidine catabolism has been shown to actively drain the cellular tetrahydrofolate pool.[23] SAM is the universal methyl donor for all cellular DNA and histone methyltransferases. Hypermethylation and presumed silencing of FTCD in FXTAS brain would be predicted to impair folate catabolism, reduce methylene-THF generation, decrease SAM synthesis, and thus reduce the overall capacity of the cell to maintain DNA methylation.

### 4.6 Myelination and White Matter Pathology

White matter disease, including T2-hyperintense lesions in the middle cerebellar peduncles and subcortical white matter, is a defining neuroimaging feature of FXTAS in males. Our data raise a candidate epigenetic mechanism for this pathology through the hypomethylation of LINGO1. LINGO1 is a co-receptor within the Nogo receptor complex that actively inhibits oligodendrocyte precursor differentiation, myelination, and axonal regeneration through activation of RhoA.[20] In the normal adult CNS, LINGO1 expression is low, permitting baseline myelination maintenance; upregulation of LINGO1 is associated with demyelinating disease, and anti-LINGO1 antibodies (opicinumab) have been investigated as remyelinating therapy in multiple sclerosis.[20] Hypomethylation-driven LINGO1 overexpression in FXTAS could impair remyelination capacity and contribute to progressive white matter deterioration. Additional hypomethylated neuronal genes, including NKAIN1 (Na+/K+ ATPase regulation), ADD2 (synaptic cytoskeleton), and COLQ (acetylcholinesterase anchoring at the neuromuscular junction), further suggest widespread disruption of neuronal membrane and synaptic function.

### 4.7 Zinc Homeostasis Dysregulation

SLC39A4 (ZIP4), encoding a high-affinity zinc importer, carried 2 hypomethylated DMPs, suggesting potential upregulation of zinc import capacity. The cerebellar-expressed zinc finger transcription factors ZIC1 and ZIC4 were also hypomethylated, with ZIC1 and ZIC4 haploinsufficiency known to cause Dandy-Walker malformation and cerebellar hypoplasia. Zinc dysregulation is increasingly recognized in neurodegenerative disease: excessive intracellular zinc contributes to mitochondrial dysfunction and neuronal death, while deficient zinc impairs synaptic signaling and antioxidant defenses. FMRP, whose levels are modestly reduced in FXTAS, is known to regulate the translation of multiple zinc transporter mRNAs.[21]

### 4.8 Non-Coding RNA Methylation

Notably, 16 of the 112 DMPs (14.3%) mapped to non-coding RNA genes, including LINC02931 (5 DMPs, hypomethylated), LINC02116 (4 DMPs, hypomethylated), LINC01623, LINC01237, LINC02525, and MIR548AA2. Long intergenic non-coding RNAs (lncRNAs) are increasingly recognized as regulators of chromatin state, transcription, and RNA processing, and their dysregulation has been implicated across multiple neurodegenerative diseases including Alzheimer’s, Parkinson’s, and the spinocerebellar ataxias.[28] The identification of methylation changes at multiple lncRNA loci raises the possibility that non-coding RNA dysregulation contributes to the FXTAS epigenetic phenotype, potentially through altered chromatin accessibility or regulation of neighboring protein-coding genes.

### 4.9 Overlap with Other Neurodegenerative Diseases

Several DMP-associated genes have established roles in other neurodegenerative conditions, suggesting shared epigenetic mechanisms across the spectrum of late-onset brain diseases. CYP2E1 hypomethylation connects FXTAS to Parkinson’s disease oxidative stress pathways.[14] RCOR1 (CoREST complex), hypermethylated in our dataset, is a transcriptional repressor whose disruption has been implicated in Huntington’s disease.[24] LINGO1 hypomethylation links FXTAS to multiple sclerosis,[20] while TRAF3 (NF-κB pathway) is implicated in neuroinflammatory processes common to multiple neurodegenerative conditions. This overlap supports the concept of a shared epigenetic landscape underlying late-onset neurodegeneration across genetically distinct disorders.

### 4.11 Limitations

Several limitations of this study should be considered when interpreting these findings. We have framed all pathway-level observations in this manuscript as exploratory and hypothesis-generating rather than confirmatory for this reason. Second, DNA methylation was profiled from bulk prefrontal cortex tissue, which is a heterogeneous mixture of neurons, astrocytes, oligodendrocytes, microglia, and endothelial cells. A single-nucleus transcriptomic data from an independent FXTAS-spectrum cohort suggest that at least some of the relevant biology (glial inflammatory gene expression) may be cell-type-specific,[34] which our bulk-tissue design cannot directly resolve. While SVA was applied to control for latent confounders, it cannot fully separate cell-type effects from disease-specific methylation changes. Third, this is a cross-sectional study of end-stage FXTAS brain tissue, and it is therefore not possible to determine the temporal relationship between methylation changes and disease onset or progression. Functional validation of the proposed epigenetic mechanisms is a key future direction.

With respect to the proteomic analysis specifically, additional caveats apply. Brain tissue is a material limitation; there could be a systematic difference in tissue handling, agonal state, or protein preservation in the brain bank repositories. We mitigated this in two ways, applying SVA to remove latent non-biological variation, and benchmarking against an independent FXTAS proteome, but residual confounding cannot be excluded. The Olink cohort was also smaller than the methylation limiting power and was restricted to a single targeted inflammation panel rather than an unbiased proteome-wide survey, so the findings reflect the inflammation-related proteome and should not be generalized to all brain proteins. NPX values represent relative rather than absolute protein quantification. The methylation and proteomic analyses were performed on overlapping but not identical sample sets, so formal per-sample integration was not performed.

The EPIC methylation array is not designed to directly predict RNA expression or protein abundance. Many EPIC probes are located in regions that don’t have a direct regulatory effect on transcription, and even when methylation affects RNA levels, protein abundance is influenced by translation, secretion, degradation, and cell-type composition. Therefore, strong one-to-one correlations between the EPIC methylation array and inflammatory proteins are not expected. Despite this limitation, the correlated DMP–protein findings are biologically informative because the associated proteins cluster around inflammatory signaling.

## 5. Conclusions

This study provides the first genome-wide DNA methylation map of FXTAS postmortem prefrontal cortex, revealing 108 significant CpG DMPs across 80 genes. These findings establish FXTAS as a disorder characterized by pervasive epigenetic reprogramming beyond the *FMR1* locus, nominating candidate gene-level disruptions in oxidative stress (CYP2E1), one-carbon metabolism (FTCD), myelination (LINGO1), zinc homeostasis (SLC39A4, ZIC1/4), synaptic function (SYT3), chromatin regulation (RCOR1, TADA2A), and — the best-supported theme in this dataset — innate immune/NF-κB signaling (TRAF3, BATF, RCOR1, MSI2).

The hypermethylation of TRAF3, the single most significant DMP in the study, together with independent downregulation of TLR3, IRAK1/4, and complement C1QA at the protein level in the same brain region, supports a coherent model of dysregulated neuroinflammatory signaling in FXTAS, evidence built from gene-level CpG replication and an orthogonal proteomic dataset. We regard the convergence of TRAF3 hypermethylation with NF-κB-pathway protein suppression as the most robust and actionable finding of this work, and the CYP2E1-DHA-maresin hypothesis (Section 4.4) as a promising but explicitly speculative avenue for future lipidomic and functional studies.

The FTCD one-carbon cycle hypothesis, if validated, would represent an important secondary amplification mechanism: epigenetic silencing of a SAM-generating enzyme could deplete the methyl donor pool and promote global hypomethylation, creating a self-perpetuating epigenetic cascade. Future studies integrating DNA methylation, transcriptomics, proteomics, and metabolomics in FXTAS brain tissue, as well as longitudinal epigenome-wide association studies in premutation carriers and direct functional validation of the candidate genes nominated here, will be essential to build upon these findings and translate them toward biomarker development and therapeutic intervention.

## Declarations

### Ethics approval and consent to participate

All brain tissue donors provided written informed consent for brain autopsy and the use of tissue and clinical information for research. The study was conducted in accordance with institutional review board (IRB) guidelines at the University of California Davis and the Icahn School of Medicine at Mount Sinai.

### Authors’ contributions

R.L. conceived and designed the study, oversaw data collection, interpreted results, and drafted the manuscript. X.L. performed bioinformatic analyses including array processing, differential methylation analysis, and pathway enrichment. D.P. contributed to study design and critical revision of the manuscript. All authors read and approved the final manuscript.

## Acknowledgments

We gratefully acknowledge the families and donors who contributed brain tissue to the UC Davis FXS/FXTAS Brain Repository led by Dr Veronica Martinez Cerdeno and the New York Psychiatric Brain Bank at Mount Sinai. We thank the staff of the Icahn School of Medicine at Mount Sinai Microarray Core Facility for array processing.

## Data availability

Raw IDAT files and processed methylation data will be deposited in the Gene Expression Omnibus (GEO) repository upon acceptance. Differentially methylated probe tables are available as supplementary data, Table 2.

